# GlomExtractor: a versatile tool for extracting glomerular patches from whole slide kidney biopsy images

**DOI:** 10.1101/2025.11.02.686103

**Authors:** Maya Maya Barbosa Silva, Justinas Besusparis, Arvydas Laurinavičius, Sabine Leh, Hrafn Weishaupt

## Abstract

The extraction of glomerular image patches is a key step prior to the training and application of glomerular classification models. Numerous algorithms for detecting and segmenting glomeruli have already been described in the literature, but standards for how to extract glomeruli from such an initial annotation remain lacking. Furthermore, the impact of different choices in extracting and preprocessing, such as cropping and scaling, are poorly understood and researched. To address this gap, the current paper introduces the GlomExtractor, a versatile tool implementing the key steps for extracting glomerular patches from whole slide images, including optional filtering of segmentations, patch extraction and scaling, and postprocessing such as background removal. By utilizing the GlomExtractor in combination with glomerular clustering experiments, the current study demonstrated how different processing strategies can impact the performance of downstream machine learning applications. We believe that the GlomExtractor and the experiments demonstrated in the current paper will help researchers in (i) the development of glomerular classification models, in (ii) advancing our understanding of the impact of patch processing on model performance, and in (iii) establishing community standards for how glomeruli should be extracted depending on downstream use.

## 1. Introduction

The histological assessment of glomeruli remains an important step in the diagnosis of non-neoplastic kidney disease. Consequently, with the expansion of the computational nephropathology field, major focus has been directed towards the development of automatic deep learning-based solutions for classifying various glomerular lesions [1–5]. Since these models typically work on glomerular image patches, successful end-to-end training and use is heavily dependent on adequate techniques for extracting such patches from the raw whole slide images (WSIs) [3–8].

However, while almost universally present in the preprocessing pipeline of glomerular classification and clustering models, there seems to be little consensus let alone any golden standard for how this extraction should be conducted. For instance, as a starting point, some studies selected patches based on a glomerular detection [4], while others relied on a glomerular segmentation [3, 5, 6]. If starting from a segmentation, different strategies for filtering, cleaning, or correcting annotations have been proposed [7, 8]. Subsequently, when extracting patches from the WSIs, studies might account for varying glomerular sizes by different scaling techniques. For instance, depending on how the patch size is selected relative to the glomerulus, extraction can either (i) preserve both the size and aspect ratio of the glomerulus [3], (ii) only the aspect ratio but not the size [5], or (iii) neither size nor aspect ratio [1, 2, 4, 6]. Finally, studies might replace any tissue outside of the target glomerulus with a contrastive color, such as black [3, 5] or white [9], while a green background has also been discussed in related contexts [10, 11]. Moreover, only few studies have investigated the impact of different pre-processing strategies, such as removing or keeping surrounding tissue [12], on downstream deep learning performances.

Furthermore, versatile tools that integrate all these various aspects for an efficient extraction of glomeruli and that could be utilized to investigate the effect of patch processing choices are still largely lacking from the literature. To address these gaps, we developed the *GlomExtractor*, a streamlined and flexible tool for extracting glomerular patches from whole slide images. Operating on segmentation annotations, the tool implements the major steps from filtering of segmentations, to patch extraction and postprocessing, while providing access to the various processing alternatives mentioned above. In doing so, the software not only provides a streamlined tool for extracting glomerular patches, but also represents a starting point for evaluating the impact of different preprocessing strategies. For instance, the current project demonstrated this utility by using the GlomExtractor to establish differently preprocessed variants across distinct glomerular datasets and ultimately evaluating the impact of the respective preprocessing alternatives in terms of clustering performance between globally sclerosed (GS) and non-globally sclerosed (nonGS) glomeruli, extending our previous work on glomerular clustering [13].

We are convinced that the tool will prove valuable in the future development of glomerular classification models as well as in further research of how the choice of pre-processing alternatives might influence their performance.

## 2. Approach

Figure 1 displays the overall workflow and main functionalities implemented in the GlomExtractor. The tool operates on raw WSIs and accompanying segmentation masks, and it’s main goal is the extraction of individual image patches centered on glomeruli (Fig. 1A). Towards this goal, it implements several functions for filtering of segmentation annotations and processing of glomerular patches (Fig. 1B). Specifically, filtering can be performed either on the image content (in terms of amount of background pixels and image blurriness) or based on the shape of the annotation (in terms of size and circularity/convexity), while processing choices include the scaling of patches and replacing the tissue surrounding each glomerulus with a contrastive color.

**Fig. 1.**
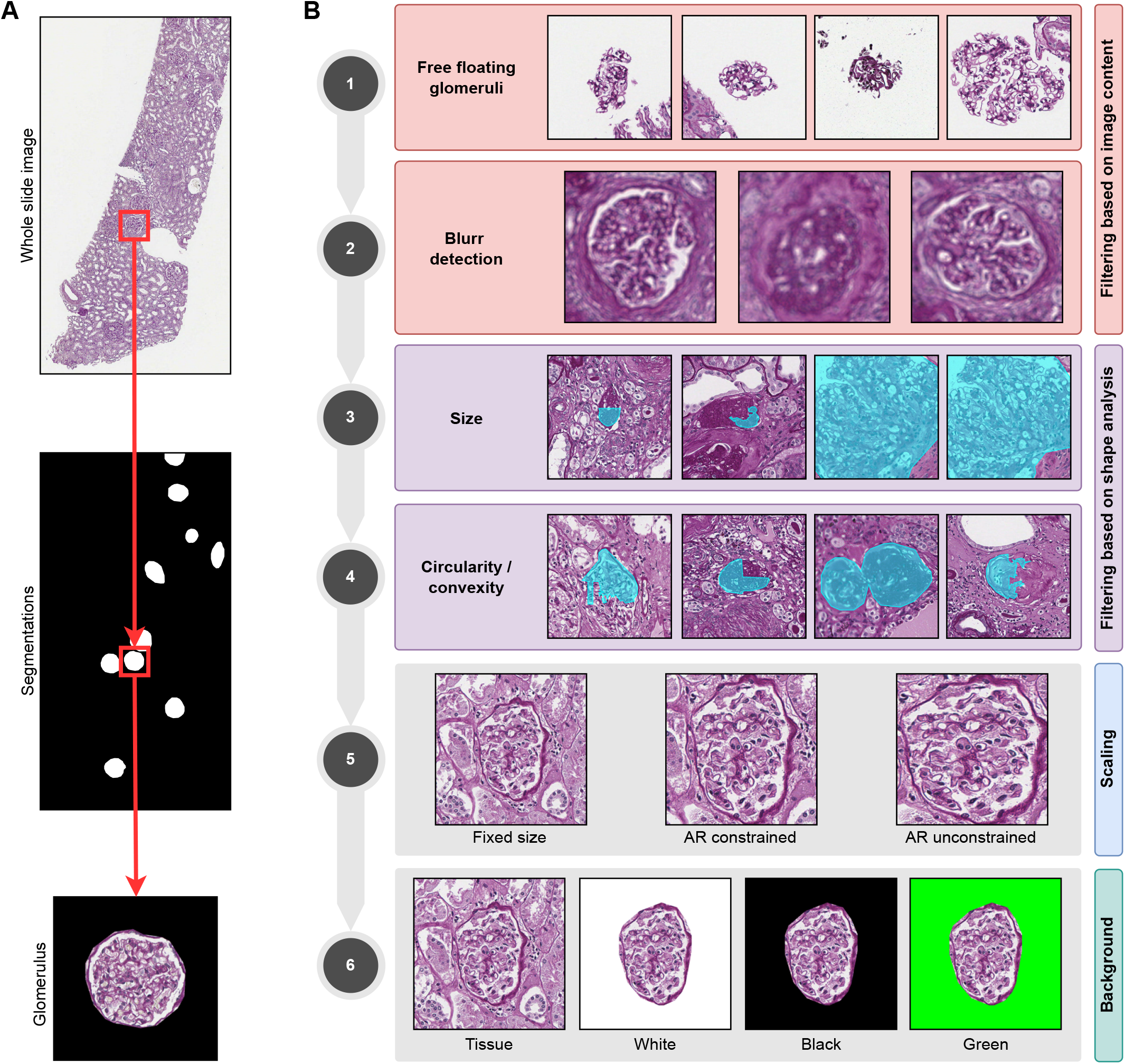
Overall workflow and functionalities implemented in the GlomExtractor. **A)** The main goal of the tool is the extraction of glomerular image patches from WSIs using accompanying segmentation annotations. **B)** The individual functions/steps implemented towards the extraction of glomerular patches: annotations can be filtered (1) to exclude glomeruli with hardly any surrounding tissue (detached from biopsy core), (2) regions with detectable blurriness, (3) exceptionally small or large annotations, or (4) non-circular/non-convex shapes; (5) patches can be extracted with different constraints on the preservation of size and aspect-ratio (AR); and (6) surrounding tissue can be replaced with a user defined color. Illustrations based on the PAS dataset.

## 3 Material and Methods

### 3.1 Data

To illustrate the functionality of the GlomExtractor, the current paper utilized a publicly available repository of kidney biopsy images and a publicly available tool for glomerular segmentation. Namely, Periodic-Acid Schiff (PAS, n=50) and Hematoxylin & Eosin (HE, n=50) WSIs were collected from the Kidney Precision Medicine Project (KPMP)[14] and segmentations were performed using the Histo-Cloud [15], as previously described [13]. To compare the effect of various scaling and cropping alternatives, the current study evaluated the clustering between globally sclerosed (GS) and non-globally sclerosed (nonGS) glomeruli in three curated and labeled datasets previously described by Weishaupt et al. [13], i.e. the Besusparis2023 (n=3993 glomeruli)[3], the Bueno2020 (n=946 glomeruli)[16] and the KPMP-PAS (n=5978 glomeruli)[14] datasets. The raw KPMP WSIs were available at 40x magnification (∼0.2527 µm/pixel), the Bueno2020 WSIs were available at 20x magnification (∼0.4936 to 0.4991 µm/pixel), while the Besusparis2023 data comprised pre-extracted patches of 1024 *×* 1024 pixels centered on glomeruli [13]. The glomerular annotations were identical to what was described in the previous paper, but patch extraction in the current study included additional options for scaling and background removal.

### 3.2 Implementation of the GlomExtractor

The GlomExtractor is implemented as a Python library with a suit of functionalities for filtering and processing of glomerular image patches, which are described in more detail below.

#### Filtering based on image content

As a first stage of preprocessing, a collection of glomerular images can be filtered based on the image content, to either remove images with extensive amounts of background areas or potentially blurry images. Specifically, for reasons of between-patch comparisons, the GlomExtractor inspects the content of a fixed-sized region (with a user-defined size) around each segmented/annotated object.

Towards the removal of images largely composed of background, the GlomExtractor measures the proportion of background pixels in the image and then allows the user to set a threshold above which images should be excluded from the dataset (Fig. 2A). The detection of background pixels utilized the `threshold_multichannel`function from the `HistomicsTK`library and is conducted on a down-scaled patch of the fixed-sized region (here 2048 *×* 2048 pixels) for the sake of computational speed. The detection of potentially blurry images relies on estimating the Laplacian variance for each image, and the user can select a threshold below which images are excluded (Fig. 2B).

**Fig. 2.**
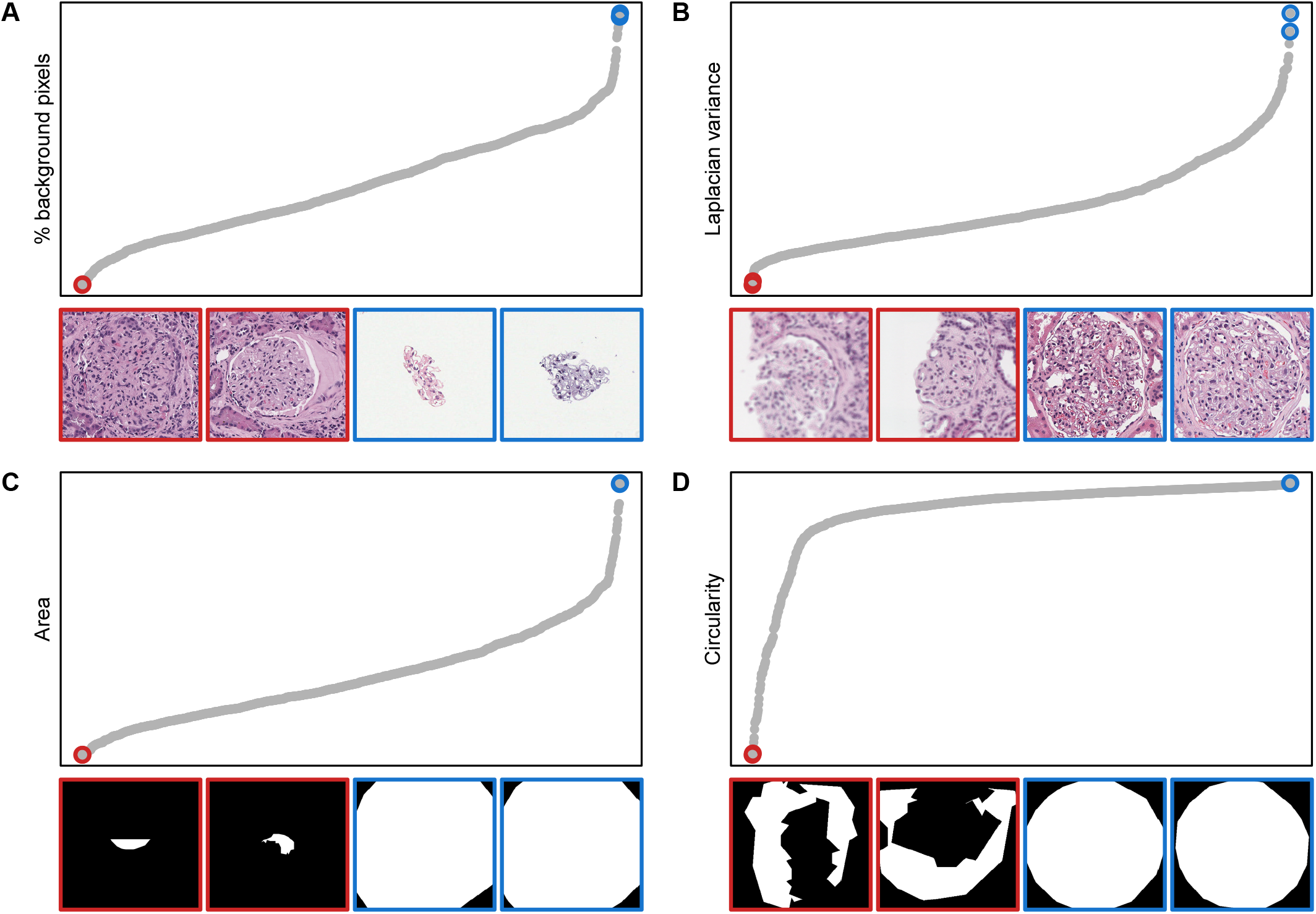
Illustration of different filtering functionalities, utilizing the the HE dataset. **A-B)** filtering based on image content, either in terms of the percentage of background pixels in a patch (A) or the Laplacian variance for detecting blurry patches (B). **C-D)** Filtering based on the segmentation shape, either in terms of the annotation area (C) or the annotation circularity (D). For each subplot (A-D), the top panel shows the measured quantity in ascending order across all patches, while the bottom panel shows the image (A-B) or segmentation masks (C-D) of the two patches with the lowest metric value (red) and two patches with the highest metric value (blue). Panels A-C show glomeruli extracted with a fixed-sized window (1024 *×* 1024), while panel D shows glomeruli extracted with dependent scaling.

#### Filtering based on shape analysis

An additional type of filtering then allows the removal of glomeruli based on their size and shape, as measured on the annotation polygon describing each segmented object. Specifically, the tool measures both the area (Fig. 2C) and the circularity (Fig. 2D). By choosing adequate thresholds, the user can then decide to remove objects with very small or large sizes (Fig. 2C), and/or a very low circularity (Fig. 2D).

#### Scaling of image patches

When extracting annotated regions into final patches, the GlomExtractor implements four different types of scaling alternatives, either based on static, fixed-sized regions around each annotation (Fig. 3A) or a more dynamic adap-tation to the bounding box around each object (Fig. 3B). For example, the user can select a certain size and either require all patches to strictly adhere to this size to ensure an identical magnification across all patches (Fig. 3C), or allow scaling-on-demand in cases where the annotation exceeds the desired fixed size (Fig. 3D). Alternatively, when basing the extracted patch on the bounding box, the user can also decide to either extract exactly the content of the bounding box, which might lead to independent/asymmetric (not preserving the aspect ratio) scaling when the image is reshaped to square dimensions (Fig. 3E), or allow for an expansion of the bounding box to facilitate dependent scaling, i.e. preserving aspect ratios, when reshaping the image region into a square patch (Fig. 3F).

**Fig. 3.**
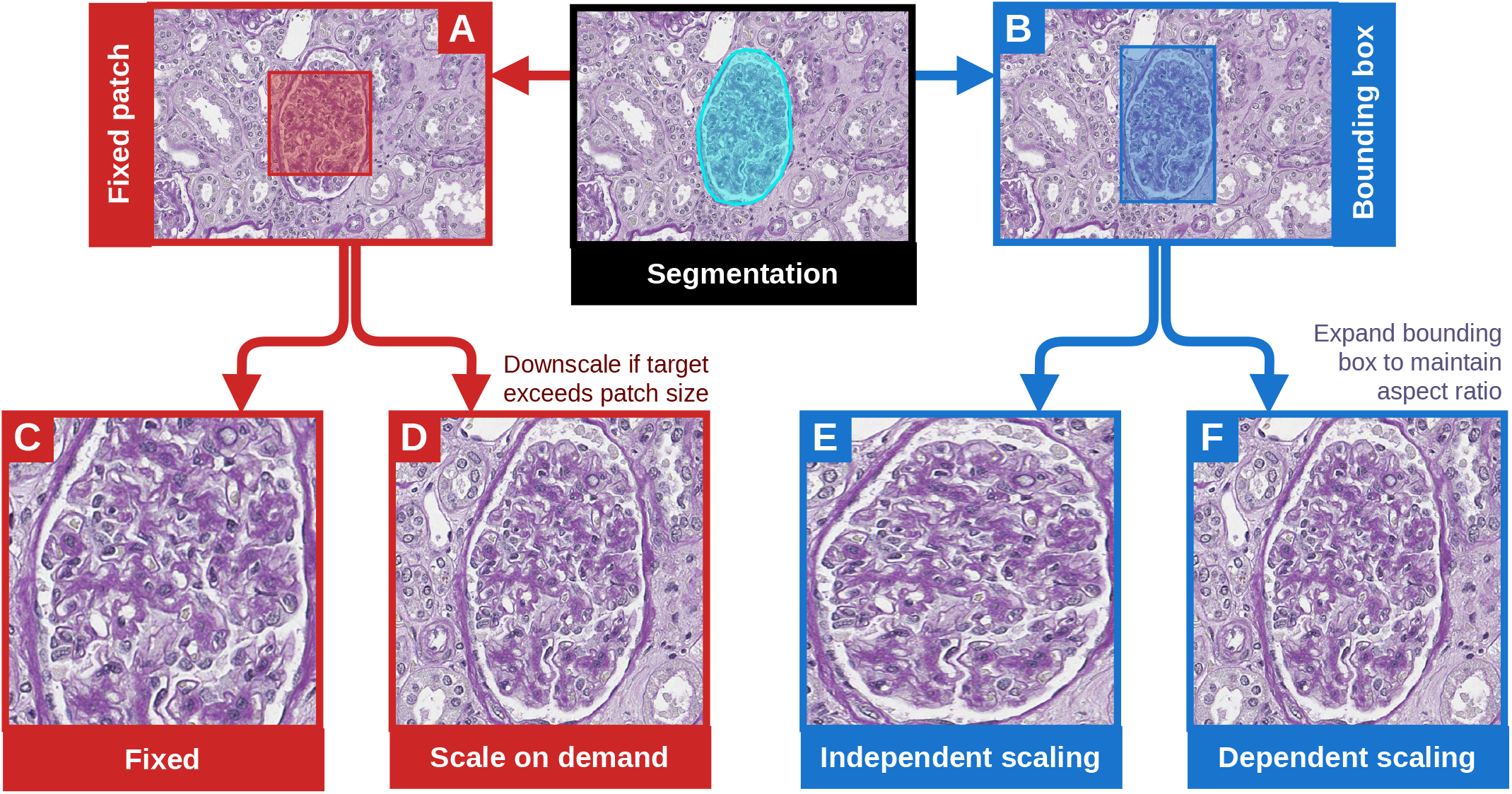
Illustration of the various scaling alternatives implemented in the GlomExtractor. **A**,**C**,**D)** Extraction of patches based on a user-defined fixed window size (A,C), including the option to downscale to fit the object inside the patch if the annotation is too large (D). **B**,**E**,**F)** Extraction of patches based on the bounding box around each object (B), either forcing the extraction of the exact bounding box which can lead to independent scaling that does not preserve the aspect ratio (E), or allowing an expansion of the bounding box to guarantee dependent, aspect-ratio preserving scaling (F).

#### Removal of non-glomerular tissue

Finally, for each patch, the tool allows the replacement of any tissue outside of the segmented object with a userdefined color, such as black, green, or white (Fig. 1). Considering that patch extraction is based on provided segmentation annotations, the GlomExtractor computes and stores a complementary segmentation mask. This mask is then used to identify and replace all non-object pixels in each patch with the desired color values.

#### 3.3 Clustering analyses

In the final stage of the current project, a set of experiments were conducted to evaluate the effect of differ-ent scaling and cropping alternatives on glomerular clustering and to demonstrate the use of the GlomExtractor in preparing datasets for such investigations. Clustering experiments were conducted as previously described [13]. Briefly, (i) patches were normalized 20 times using the different reference images selected for the respective histological stain in Weishaupt et al. [13], (ii) features were extracted using the MobileNet [17], (iii) clustering was performed using the Leiden algorithm [18], following the strategy described by Weishaupt et al. [13], and cluster performance was evaluated using the Adjusted Rand Index [19]. However, in addition to the single patch extraction conducted in our previous paper, i.e. independent scaling and replacing non-glomerular tissue with black color (independent-black), five additional combinations of scaling and background colors were employed prior to the clustering analyses: (i) fixed-size (1024 *×* 1024 pixels for Bueno2020 and Besusparis2023, and 1780 *×* 1780 pixels (width of largest glomerulus) for KPMP-PAS) patch extraction with black background, (ii) dependent scaling with black background, (iii) independent scaling without removing the surrounding tissue, and independent scaling with (iv) green or (v) white background. These datasets allowed to compare the difference in clustering performance between the various scaling techniques (independent/dependent/fixed), between keeping or removing surrounding tissue, and between different colors use to replace the non-glomerular tissue. Performances were always compared with respect to the independent-black dataset, which was the pre-processing alternative employed in the original paper [13], and statistical significance of differences was evaluated using a two-tailed, paired Wilcoxon signed-rank test using the `wilcox.test`function in R, i.e. each of the final six dataset resulted in 20 clustering performance values, one per stain normalization reference, with pairing between datasets then based on the stain reference. The p-values within each dataset were adjusted for multiple testing via False Discovery Rate (FDR) correction utilizing the `p.adjust`function in R, and a comparison was considered significant if the resulting adjusted p-value was *q <* 0.01. Visualizations were performed in R.

### 3.4 Code availability and usability

The GlomExtractor is developed in Python and is dependent on common libraries, most importantly but not only, *OpenSlide,TiffFile* and *Shapely*. The tool is available as a command-line interface designed to receive as input the directory containing both original WSIs and their respective annotations. The user also has different options to select based on their desired filtering and processing approach. A more detailed description of the user inputs and their default values can be seen on the toolkit’s GitHub page. All filtered-out images picked by the tool can be saved to a separate folder for manual inspection and post-correction when the tool is used in its *debug* function.

The source code and documentation of the GlomExtractor is made freely available at https://github.com/patologiivest/GlomExtractor.

## 4. Results

To demonstrate the use of the GlomExtractor, the current study employed it to evaluate the effect of patch processing choices on glomerular clustering between globally sclerosed (GS) and non-globally sclerosed (nonGS) glomeruli [13]. Specifically, obtaining three labeled glomerular datasets [3, 14, 20] and a general clustering strategy from our previous publication [13], the performance effects of scaling (Fig. 4A), removal of surrounding tissue (Fig. 4B), and the choice of background replacement color (Fig. 4C) were investigated.

**Fig. 4.**
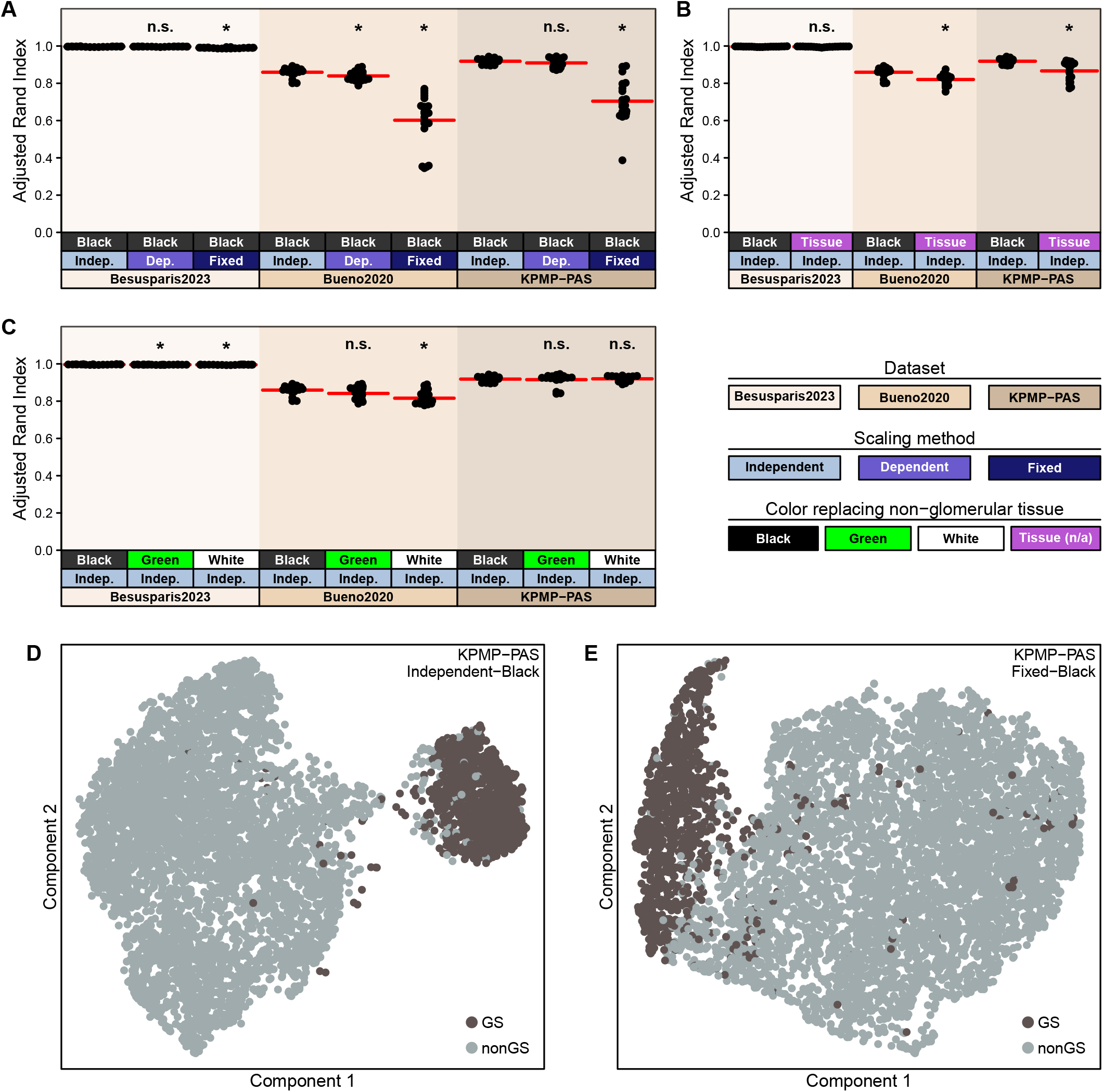
Comparing the effects of patch preprocessing choices on clustering performance. **A-C)** Strip charts displaying clustering accuracy in terms of the adjusted Rand Index in three different datasets (Besusparis2023, Bueno2020, KPMP-PAS), comparing either different scaling approaches (A), removal/non-removal of non-glomerular background (B), and different background replacement colors (C). Each strip contains 20 values, one for each of the 20 stain-normalized versions of the respective dataset, with the red lines indicating the corresponding mean value. *: *q <* 0.01 (paired, two-tailed Wilcoxon signed rank test, FDR-adjusted within each dataset). **D-E)** UMAP embedding of one stain-normalized KPMP-PAS variant extracted either with independent scaling (D) or as fixed-sized windows (E). Light and dark colored dots represent nonGS and GS glomerular patches.

As previously reported [13], the Besusparis2023 dataset achieved almost perfect clustering in the original dataset (independent scaling with black background), and only very minor absolute changes were observed with the other preprocessing strategies (Fig. 4), likely because the separation between the classes was too pronounced to be substantially affected. Accordingly, downstream performance comparisons were based more on the Bueno2020 and KPMP-PAS datasets, which exhibited more pronounced absolute performance differences (Fig. 4).

Specifically, in both aforementioned datasets, patch extraction using fixed-sized windows (Fig. 4A) or retaining non-glomerular tissue (Fig. 4B) resulted in significantly reduced clustering performances. Importantly, with fixed-sized patch extraction, some clustering runs essentially failed, resulting in adjusted Rand index (ARI) values below 0.4 (Fig. 4A), likely because the resulting feature embedding did not lead to a sufficient separation between the classes (Fig. 4D-E). With respect to the comparison between independent and dependent scaling (Fig. 4A) or between the different background replacement colors (Fig. 4C), some significant effects were observed, but the patterns were not consistent across datasets.

## 5. Discussion

The extraction and preprocessing of patches is a fundamental step in most end-to-end glomerular classification models. However, few efforts have addressed (i) the establishment or evaluation of consensus strategies for how such extractions should best be conducted or (ii) the development of freely available software tools allowing flexible and versatile solutions for all aspects related to patch extraction and processing. As a solution to these challenges, the current study presents the GlomExtractor, an opensource toolkit for streamlined processing of glomerular image patches, including options for filtering, extracting, scaling, and cropping glomeruli from WSIs. We believe that this software can substantially aid researchers in generating glomerular datasets, and, importantly, support further studies into the effects of glomerular processing choices.

The clustering experiments conducted during the current study indicate that the choice of patch extraction and processing strategy can indeed have pronounced effects on downstream clustering performance. For instance, the strongest and most consistent performance drops among the Bueno2020 and KPMP-PAS datasets were observed when utilizing the fixed-sized window extraction method or opting for the retention of non-glomerular tissue. An explanation for the former scenario might be that feature embedding and clustering are influenced by features related to glomerular area and by largely varying amounts of input in terms of the ratio of pixels within and outside of the glomerulus, since a “fixed-sized” extraction, by design, preserves glomerular sizes. The latter observation might be due to the influences of heterogeneous features extracted from the non-glomerular tissue areas. A similar effect was observed in a recent study by Trejo et al. [12], where the performance of their glomerular classification model worsened when trained on glomerular patches that maintained non-glomerular tissue. On the other hand, it is still unclear, whether the area surrounding the glomerular capsule might include relevant information for e.g. the classification of some morphological lesions. Furthermore, although not investigated in the current study, a combination of fixed-sized patch extraction and retention of background might face even further problems in clustering, especially in cases were the patch contains additional glomeruli, beyond the original target, that exhibit different morphological classes.

Evaluations of the Besusparis2023 dataset were less conclusive, since the dataset revealed an almost perfect separation between the two classes that was not very susceptible to variations in patch processing. The excellent separation was attributed to the fact that the dataset appeared very clean, likely because it was highly curated [13] and perhaps also because of some specific properties of the utilized modified Picrosirius red stain. Consequently, the impact of patch processing might also depend on how clean the dataset is and/or how distinct the target classes are.

Altogether, while showcasing the usability of the GlomExtractor, the conducted experiments and obtained findings serve only as a first initiative into revealing the impact of patch extraction choices on downstream machine learning performances. Further experiments will clearly be needed to fully investigate the extent of these effects on different applications, such as classification or clustering of other glomerular lesions, and in order to establish consensus recommendations for preprocessing strategies.

Importantly, beyond providing a highly versatile toolkit for the processing of glomerular patches, the GlomExtractor can in principle be applied to any kind of object as long as the WSI is provided with appropriate segmentation annotations. The tool also exhibits several avenues for further improvement. For instance, while, at present, it only includes texture-based analysis based on the estimation of background pixels and Laplacian filters, the inclusion of other types of texture or intensity features [21] might add further value. Similarly, while the Laplacian filter might help in the detection of some blurred images, no other artifact detection techniques are currently implemented. Nevertheless, it would be possible to run separate artifact detection tools, such as the HistoQC [22], and exclude suspicious WSI regions prior to patch extraction. Finally, to increase the usability of the tool, it would be beneficial to explore options to facilitate (i) graphical user interfaces easing the filtering and selection of glomeruli during dataset generation and/or (ii) direct integration into end-to-end deep learning pipelines.

## 6. Conclusion

In summary, the GlomExtractor is a publicly available and highly versatile tool for glomerular patch handling, including various options for annotation filtering, patch extraction, and patch postprocessing. As illustrated in the current study, the GlomExtractor can serve as an efficient starting point to investigate the impact of glomerular patch processing on downstream machine learning performance, the possible effects of which were clearly demonstrated in the showcased experiments. We hope that the GlomExtractor will aid researchers in generation of glomerular datasets and in furthering the community understanding and standards related to patch processing choices. Future improvements of the software could focus on (i) the inclusion of additional feature extraction and filtering methods, (ii) additional artifact detection functionalities, and (iii) improving its usability through graphical user interfaces and/or integration into end-to-end DL training frameworks.

## Acknowledgments

The results here are in part based upon data generated by the Kidney Precision Medicine Project (https://www.kpmp.org). The project was funded by The Western Norway Health Authority (strategic research fund F-12563 and open project fund F-13120).

## Author contributions

*Conceptualization*: H.W.; *Data curation*: H.W., J.B.,; *Formal analysis*: H.W.; *Funding acquisition*: S.L.; *Investigation*: H.W.; *Methodology*: H.W.; *Project administration*: H.W.; *Resources*: A.L., S.L.; *Software*: H.W., M.M.B.S; *Supervision*: H.W.; *Validation*: H.W.; *Visualization*: H.W.; *Writing – original draft*: H.W., M.M.B.S.; *Writing – review and editing*: H.W., M.M.B.S., J.B., A.L., S.L.

## Notes

### Competing Interest Statement

The authors have declared no competing interest.

